# Understanding narwhal diving behaviour using Hidden Markov Models with dependent state distributions and long range dependence

**DOI:** 10.1101/388835

**Authors:** Manh Cuong Ngôe, Mads Peter Heide-Jørgensen, Susanne Ditlevsen

**Affiliations:** Greenland Institute of Natural Resources, c/o Greenland Representation, Strandgade 91,2, 1401 Copenhagen K, Denmark; Department of Mathematical Sciences, University of Copenhagen, Universitetsparken 5, 2100 Copenhagen, Denmark

## Abstract

Diving behaviour of narwhals is still largely unknown. We build three-state Hidden Markov models (HMM) to describe the diving behaviour of a narwhal and fit the models to a three-dimensional response vector of maximum dive depth, duration of dives and post-dive surface time of 8,609 dives measured in East Greenland over 83 days, an extraordinarily long and rich data set. In particular, diurnal patterns in diving behaviour for a marine mammal is being inferred, by using periodic B-splines with boundary knots in 0 and 24 hours. Several HMMs with covariates are used to characterize dive patterns. Narwhal diving patterns have not been analysed like this before, but in studies of other whale species, response variables have been assumed independent. We extend the existing models to allow for dependence between state distributions, and show that the dependence has an impact on the conclusions drawn about the diving behaviour. It is thus paramount to relax this strong and biologically unrealistic assumption to obtain trustworthy inferences.

**Author summary:** Narwhals live in pristine environments. However, the increase in average temperatures in the Arctic and the concomitant loss of summer sea ice, as well as increased human activities, such as ship traffic and mineral exploration leading to increased noise pollution, are changing the environment, and therefore probably also the behavior and well-being of the narwhal. Here, we use probabilistic models to unravel the diving and feeding behavior of a male narwhal, tagged in East Greenland in 2013, and followed for nearly two months. The goal is to gain knowledge of the whales’ normal behavior, to be able to later detect possible changes in behavior due to climatic changes and human influences. We find that the narwhal uses around two thirds of its time searching for food, it typically feeds during deep dives (more than 350 m), and it can have extended periods, up to 3 days, without feeding activity.

## 1 Introduction

The narwhal (*Monodon monoceros*) is a long-lived toothed whale that primarily inhabit cold waters of the Atlantic sector of the Arctic. The largest abundances are found in East and West Greenland and in the Canadian High Arctic. Physical maturity is reached at an age of ~ 30 years and at a body length and mass of 4 – 5 *m* and 1,500-1,800 kg [8]. The narwhal belong to the group of the deepest diving cetaceans with the maximum exceeding 1, 800 *m* [10]. Their diet consists of few prey species including Greenland halibut (*Reinhardtius hippoglossoides*), polar cod (*Boreogadus saida*), capelin (*Ammodytes villosus*) and squids (*Gonatus sp*.) [9,13]. Predation on Greenland halibut constitutes a potential competition with fisheries, especially in winter, at the same time as narwhal skin and meat as well as their valuable tusks for centuries have been important hunting products for Inuit cultures in Greenland and northern Canada. Anthropogenic factors like underwater noise are also a concern for a species that, with decreasing sea ice coverage, is increasingly exposed to underwater noise from shipping and seismic exploration [21]. It is therefore important to understand and quantitatively describe the diving activities of narwhals, by robust statistical methods, and assess how these are changed due to potential conflicts with anthropogenic activities, and to ensure the long-term conservation of one of the most specialized species in the North Atlantic.

The first step is to understand the diving patterns of narwhals under natural conditions, which we address in this study. Diving behaviour is however cryptic since it includes both physiological constraints, energetic demands and habitat and environmental regimes. Modelling of the observed diving behaviour is one way of gaining insight to the overall diving patterns, and changes in model parameters is a way to compare and estimate quantitatively impacts from anthropogenic activities (see [4,5]).

In this study we apply multivariate Hidden Markov Models (HMMs) with covariates [22], to describe the diving dynamics in the vertical dimension of an individual narwhal. These types of models for similar diving data of Blainville’s beaked whales (*Mesoplodon densirostris*) were first introduced in [15]. A HMM assumes an underlying unobserved process, which governs the dynamics of the observed variables. The assumption is that the observed behaviour in a dive will depend on the present state, and introduces autocorrelation in the model [22], in agreement with the observed data, where the narwhal tends to repeat the same type of dives in a series before switching to another type of dives. Similar to the blue whales (*Balaenoptera musculus*) data analysed in [5] and the short-finned pilot whales (*Globicephala macrorynchus*) data analysed in [19], our narwhal data suggest three distinct states: state 1 covers the shortest and shallowest dives which we interpret as near-surface travelling, state 2 covers the longer and deeper dives which we interpret as feeding states, and state 3 covers the deepest and longest dives which we interpret as intense deep feeding behaviours. Pohle et al [18] recommended against using more than four states in biological modelling like this, in order to avoid the complexity of the correspondence between states of the model and the biological phenomenon. DeRuiter et al [5] suggested three states for their data, even if a formal model selection procedure would point to a more complex model, because models with more underlying states might obscure patterns in the data and provide less insight in the underlying biological process, even if they might perform better in terms of forecasting. Biological knowledge should guide the choice of number of states. They also argue that model misspecifications, such as too inflexible state dependent distributions, variations over time, missing covariate information or outliers might cause model selection criteria to favour models with more complex structures than warranted. Therefore, we choose the three-states HMM model.

These HMMs have shown powerful for modelling animal movement by taking into account the correlation over time between different movement patterns, with an extensive research in two horizontal dimensions (see, e.g., [14,16,17]), and recently, in one vertical dimension [5,15], possibly including further information on vertical movements. In this study, we use vertical depth data, and the three response variables are the maximum depth reached in a dive, the duration of a dive, and the post-dive surface time before initiating a new dive.

The data set covers 1,995 hours (~ 83 days) and is extraordinarily long, and thus provides a unique opportunity to obtain detailed information on diving behaviour. An example of the data is shown in Fig 1. Such data are usually only on the order of a couple of days or less, for example, the time series of short-finned pilot whales analysed in [19] cover up to 18 hours and 64 dives, whereas the time series of blue whales analysed in [5] cover up to 6 hours and 67 dives, and [14] analyses 79 hours of a single Blainville’s beaked whale. However, here we only have data from a single narwhal limiting the generalizability of the analysis.

**Fig 1.**
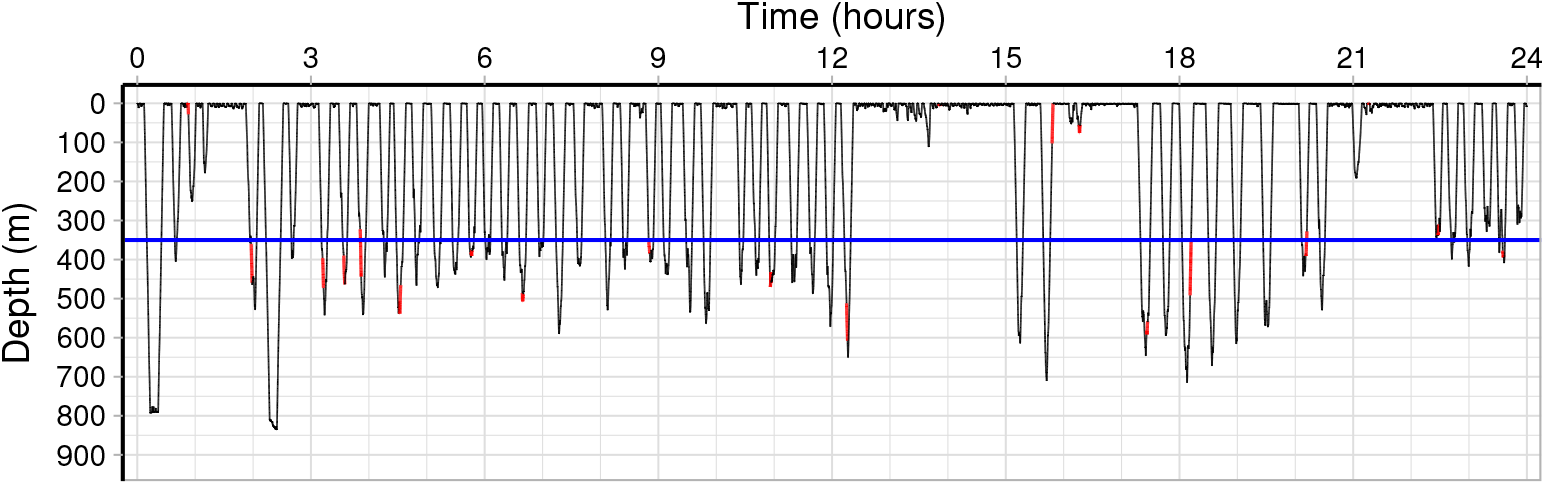
Diving data. Representative part of the narwhal diving data, covering 24 hours of dives on August 15th 2013. The red parts are where a lower temperature in the stomach has been registered, indicating that the narwhal has swallowed a prey. The blue line indicates a depth of 350*m*, the threshold for a *deep dive* used in the definition of the covariates.

In all previous studies, *contemporaneous conditional independence* was assumed, meaning that the state dependent processes are independent given the underlying state. This is a strong and biologically unrealistic assumption, since deeper dives will typically take longer, also when conditioning the dive to be either shallow, medium or deep. A positive correlation is still expected, beyond the correlation implied by the hidden states. DeRuiter et al [5] argued for the assumption of conditional independence because unless a multivariate normal distribution can be assumed, there is usually no simple candidate multivariate distribution to specify the correlation structure. This is partly due to some of their response variables being discrete. In this study, we will relax the assumption of conditional independence, taking advantage of the continuity of the response variables. They are all restricted to be positive and with right skewed distributions. Previous studies have therefore used conditionally independent gamma distributions for these variables. Here, we will assume dependent log-normal distributions, such that their log-transforms follow a multivariate normal distribution. We only assume dependence between maximum depth and dive duration, which has a high correlation in the full data set (0.864), as well as high correlations within the three estimated states for the chosen model (0.538, 0.777 and 0.458, respectively), whereas we assume both of these variables independent of post-dive surface time, since the correlations in the full data set were low (0.046 and 0.042, respectively). We also do the analysis with the standard choice of the gamma distributions, and compare the results.

Covariates were included in [5,15,16], appearing in the transition probabilities between hidden states, whereas no covariates were included in [19]. Here we include covariates in all elements of the transition matrix, trying out different covariate process models and select the optimal model by the Akaike Information Criterion (AIC). We consider two covariates related to the recent deep dives performed by the narwhal.

Dives can reach > 1, 800 *m*, and deeper dives are usually related to intense feeding [10]. We define a *deep dive* as a dive to a depth of at least 350 *m*. Note that this definition is only used to define the covariates, and is not related to the decoding of states. One covariate is the time passed since the last deep dive, which was also used in [15]. The hypothesis is that the longer the time passed since last deep dive, the higher the narwhal’s propensity for initiating a deep dive will be. Another covariate counts the number of consecutive deep dives that the narwhal has performed. The hypothesis is that the more dives in a row and more time spent at great depths, the higher the narwhal’s propensity for changing diving pattern to shallower depth or near-surface travelling. By introducing such history dependent covariates, the model allows a longer dependence structure than the one implied by the Markov property. These models with dependencies between observables caused by the underlying state, as well as including feedback from the observed process, were introduced in [15] to model Blainville’s beaked whale. The last covariate is time of day at initiation of the dive, modelled by a periodic B-spline with boundary knots in 0 and 24 hours. Diurnal effects on marine mammal diving patterns are difficult to estimate in this type of models because the time series are typically too short, but this data set is extraordinarily long, making this inference possible. However, here we only analyse a single whale, and results might not generalize.

## 2 Materials & Methods

We analyse the time series of depth measurements of a mature male narwhal (420 cm, estimated mass 950 kg) tagged in East Greenland from August 13th until November 6th 2013. Permission for capturing, handling, and tagging of narwhals was provided by the Government of Greenland (Case ID 2010–035453, document number 429 926). The tag (Mk10 time-depth recorder from Wildlife Computers, Redmond, WA, USA) was attached to the whale and retrieved one year later with 1994.83 hours of dive data (approximately 83 days and 2 hours), see [12]. In this time interval the narwhal performed 8,609 dives to depths of at least 20 *m*. Depth was measured every second at a resolution of 0.5*m*, and summarized in three variables within each dive to describe the behaviour: dive duration, maximum depth and post-dive surface time, as also used in [5,19]. Here we define a dive as one in which the whale reach a depth of at least 20 *m*, otherwise it is considered time spent at the surface. This threshold was chosen in order to avoid creating too many shallow dives near the surface, see [1]. The surface and dive durations also enter in the model as part of the covariate counting the time since last deep dive. The total time the whale spent at the surface was 37 days and 5.6 hours, i.e., the narwhal spent 44.8% of its time at the surface.

The first week of tagging, the narwhal also had the temperature of the stomach measured, see [11]. A temperature drop indicates that a prey has entered the stomach. Fig 1 shows an example part of the data, and the red parts indicate temperature drops. These typically happen during deep dives, and support the assumption that deep dives are related to foraging.

In this study, the observed response variable, denoted by ***X**_t_*, is three-dimensional, describing the diving behaviour related to each dive, where *t* indicates the dive number, *t* = 1, 2,…, *T*, with *T* = 8, 609 dives. The first response variable, *X*_1,*t*_, is the maximum depth reached in dive number *t*, and takes real positive values between 20 and 910.5 *m*. The second response variable, *X*_2,*t*_, is the duration of dive number *t*, and takes real positive values between 33 seconds and 28 minutes. The third response variable, *X*_3,*t*_, is the time spent at the surface after the dive before initiating a new dive, and takes real positive values between 1 second and 209.7 minutes. We assume that the diving behaviour depends on an underlying unobserved process, which we denote by *C_t_, t* =1, 2,…, with a number *m* of unobserved behavioural states, *C_t_* ∈ {1,…, *m*}, which govern the dynamics of the observed variables. The assumption is that the distributions of the observed maximum depth, duration of dive, and post-dive duration of dive number *t* depend on the state. We choose the three-states HMM model, *m* = 3, and identify the three states with three different diving patterns in the water column; near-surface travelling, medium depth dives, and deep dives probably related to feeding.

### 2.1 Hidden Markov Model

An *m*-dimensional hidden Markov model assumes that the distribution of the *p*-dimensional response vector ***X**_t_* depends on a hidden state *C_t_*, where {*C_t_* : *t* = 1, 2,… } is an unobserved underlying process satisfying the Markov property:

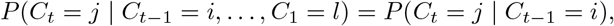

where *C_t_* ∈ {1,…, *m*} for *t* = 2, 3,…. Denote the state transition probabilities for *m* = 3 at time *t* by *ω_ij_* (*t*), *i, j* = 1, 2, 3, where *ω_ij_* (*t*) = *P*(*C*_*t*+1_ = *j* | *C_t_* = *i*). The transition probability matrix Ω(*t*) is then

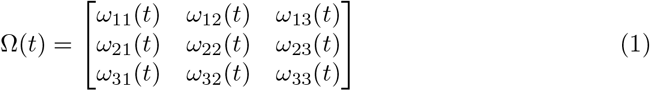

where *ω_ij_* (*t*) ≥ 0 and 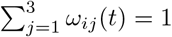. Here, we let *ω_ij_* (*t*) depend on *t* to allow time varying covariates to affect the transition probabilities, see Section 2.3. The distribution of ***X**_t_* is conditionally independent of everything else given *C_t_*:

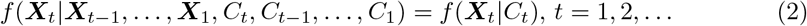

where *f* denotes a probability density function, i.e., the distribution of ***X**_t_* depends only on the current state *C_t_* and not on previous states or observations. The model is illustrated in Fig 2.

**Fig. 2.**
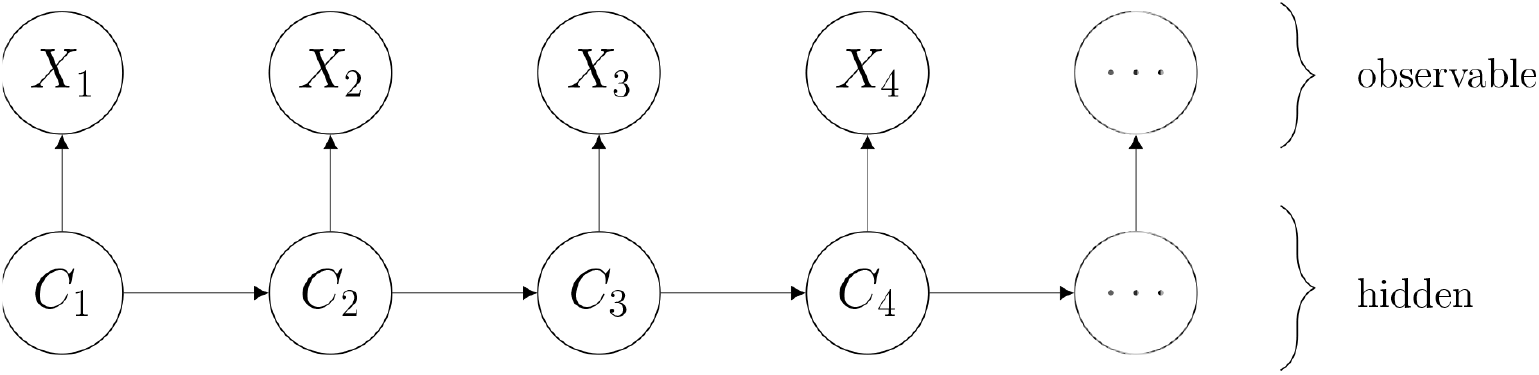
Hidden Markov Model. The hidden states *C_t_* represent behavioural states that influence the distribution of the observed variables *X_t_*.

### 2.2 State dependent distributions

The state-dependent distributions are the probability density functions of ***X**_t_* associated with state *i*. Under the *contemporaneous conditional independence* assumption, the *p* different components of the response vector ***X**_t_* are assumed independent given the hidden state, and the probability density can be decomposed as

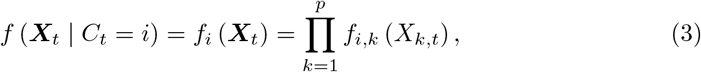

where *X_k,t_* is the *k*th observed component of ***X**_t_*. Here we have *p* = 3, the components being maximum depth (*MD*), duration of dives (*DT*), and post-dive duration (*PD*). Thus, ***X**_t_* = (*X_MD,t_*, *X_DT,t_*, *X_PD,t_*)^*T*^, where ^*T*^ denotes transposition. *Contemporaneous conditional independence* implies that the state dependent processes *X_MD,t_,X_DT,t_* and *X_PD,t_* are independent given the underlying state *C_t_*. This assumption has been used in [5] and [19] because in general, there is no simple way to address the correlation between variables within states, and the dependence induced by the Markov chain is often sufficient to fit the data. However, in this paper, we will relax this assumption, and let *f_i_* be a joint distribution function, allowing for dependent coordinates, which for our data turn out to improve the fit considerably.

All three response variables are positive right-skewed variables, so natural candidates for *f_i,k_* are gamma distributions, as used in [5] and [19], or log-normal distributions, i.e., the logarithm of the response variables follow a 3-dimensional normal distribution. Here, we will try three different distributions. The first candidate is independent gamma distributions, to compare with the usual approach. The gamma distribution is parametrized by shape parameter *μ* and scale parameter *σ*, with mean *μσ* and variance *μσ*^2^, and the state dependent probability density functions are given by

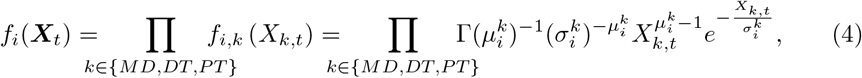

for *i* = 1, 2, 3.

We will also assume both independent and correlated log-normal distributions, such that log ***X**_t_* is multivariate normal, where log ***X**_t_* = (log *X_MD,t_*, log *X_DT,t_*, log *X_PT,t_*)^*T*^, taking advantage of the computational convenience of the normal distribution. The log-normal distribution is parametrized by log-mean *μ* and log-variance *σ*^2^. Thus, given *C_t_* = *i* and *k*, the mean and variance of log ***X**_k,t_* is 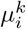 and 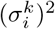, and the mean and variance of ***X**_k,t_* is 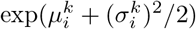 and 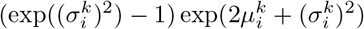. The log-correlation is denoted by *ρ_i_*. The correlation between the first two components is 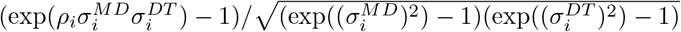, where 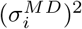 and 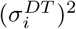 are the log-variances of maximum depth and dive duration, respectively. The correlation is approximately equal to the log-correlation *ρ_i_* when 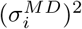 and 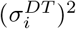 are small. We assume post-dive duration independent of the other two response variables, based on the low marginal correlation in the data. Thus, the state dependent probability density functions are given by

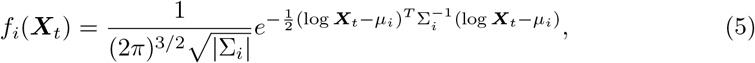

where | · | denotes the determinant of a matrix, 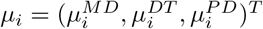,

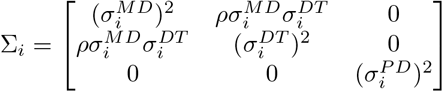

and *ρ* = 0 in the independent case.

### 2.3 Covariates

To allow for a longer memory in the model beyond the autocorrelation induced by the hidden process, we incorporate feedback mechanisms by letting the state transition probabilities depend on the history. We consider two covariates related to the recent deep dives performed by the narwhal. One covariate is the continuous variable *τ_t_*, defined as time passed since the last deep dive before dive number *t*, where a *deep dive* is defined as a dive to a depth of at least 350 *m*. Note that this definition is only used to define the covariates, and is not related to the decoding of states. Thus, the algorithm might decode dives of lesser depth to state 3, and deeper dives not to be in state 3. The value chosen is somewhat arbitrary, and we tried different values between 250 and 450 *m*, without much effect on the results. The other covariate is the discrete variable *d_t_* taking non-negative integer values, counting the number of consecutive deep dives that the narwhal has performed before dive number *t*. Finally, we consider the covariate of the hour at which the dive is initiated. More specifically, we define the covariate processes 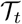, the time since the last deep dive, *D_t_*, the number of consecutive deep dives up to dive number *t*, and *H_t_*, the hour of initiation of dive *t*, and denote the measured covariates by *τ_t_, d_t_* and *h_t_*. Thus, the short term memory is modelled by the hidden states, and the long term memory is modelled by modulation of the transition probabilities as a function of past dynamics. The model is illustrated in Fig 3, and Fig 4 illustrates the response variables and the three covariates for 60 consecutive dives.

**Fig. 3.**
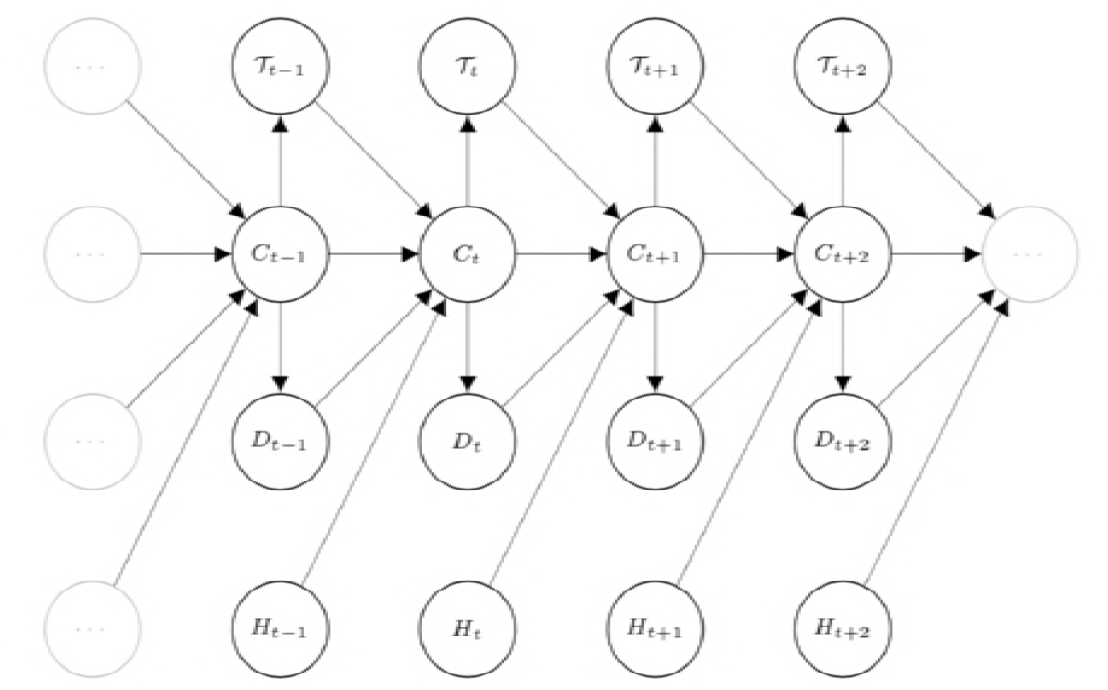
Hidden Markov Model with feedback processes. The distribution of the transitions between hidden states *C_t_* depends on the observed covariate processes 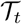, *D_t_* and *H_t_*.

**Fig. 4.**
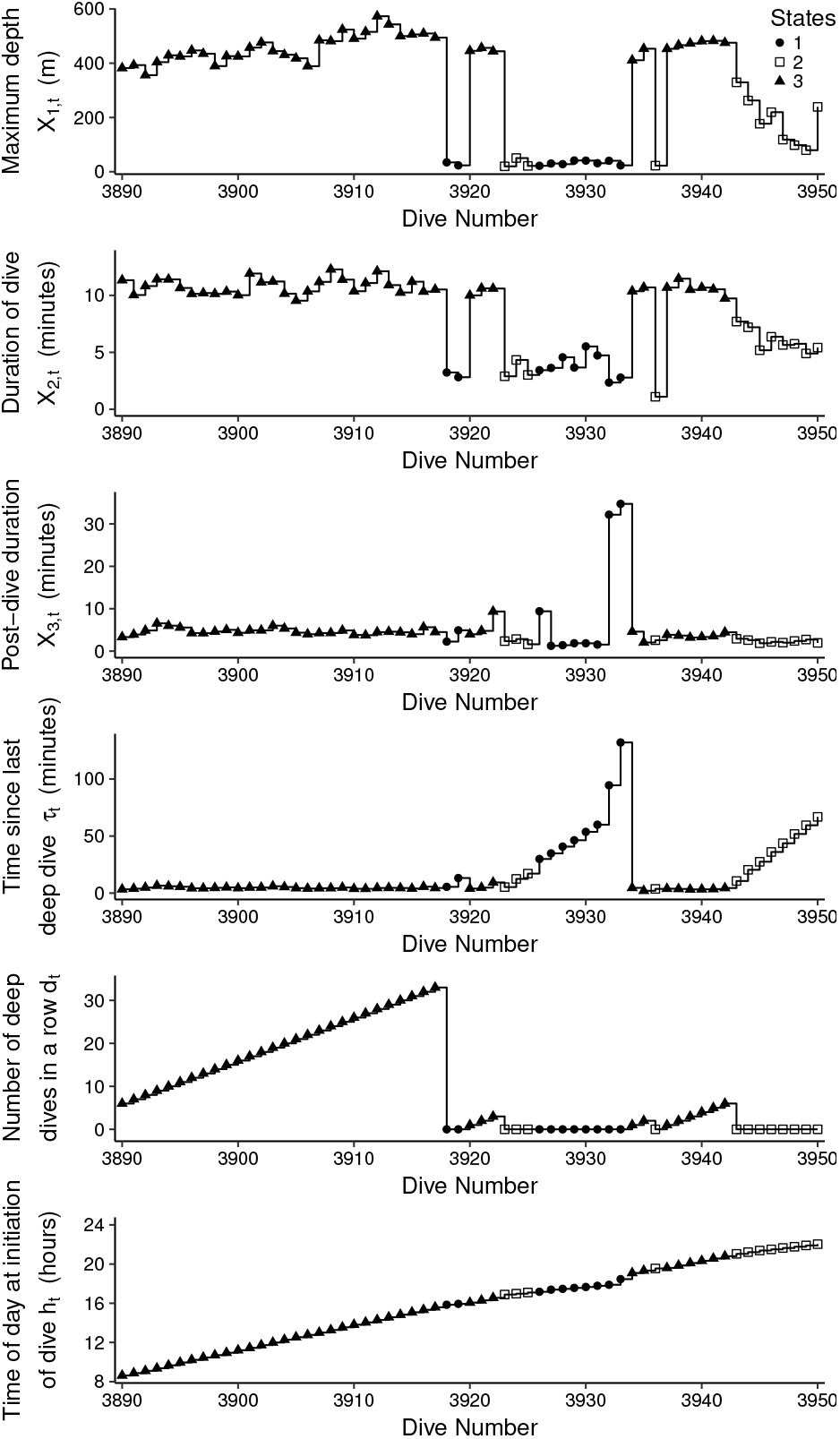
Response variables and covariate processes. Time series plot of maximum depth, duration of dive, and post-dive duration from dive number 3890 to 3950 and the covariate processes counting the time since last deep dive, number of deep dives in a row, and the hour at initiation of dive. The symbols indicate the decoded hidden states from a model fitted to a dependent log-normal distribution (Model 1). State 1 (circles) represents near-surface travelling, State 2 (squares) represents a feeding state of medium long and deep dives, and State 3 (triangles) represents an intense deep feeding state of the deepest and longest dives.

The covariates enter the transition probabilities *ω_ij_* (*t*) = *ω_ij_* (*η_ij_* (*t*)) in Eq. (1) through a *predictor, η_ij_* (*t*), see Eq. (7) below. We consider several models. If there are no covariates for a given predictor, then *η_ij_* (*t*) = *η_ij_* does not depend on *t*. Table 1 lists the different models, where *α_ij_, β_ij_, γ_ij_, δ_ij_, θ_ij_*, and *ζ_ij_* are real parameters. We tried more models, but only include these for illustration. Covariates *d_t_* and *τ_t_* were incorporated as natural cubic splines with three degrees of freedom. The covariate *d_t_* counts number of deep dives in a row, and is therefore around 0 when not in state 3. This covariate therefore carries no information unless in state 3, and only enters in *η*_31_ and *η*_32_. Likewise, *τ_t_* is expected to be around 0 when in state 3, and therefore only enters *η_ij_* for *i* = 1 or 2. The effect of time of day is modelled by a periodic B-spline with three degrees of freedom, with boundary knots in 0 and 24 hours.

**Table 1.**
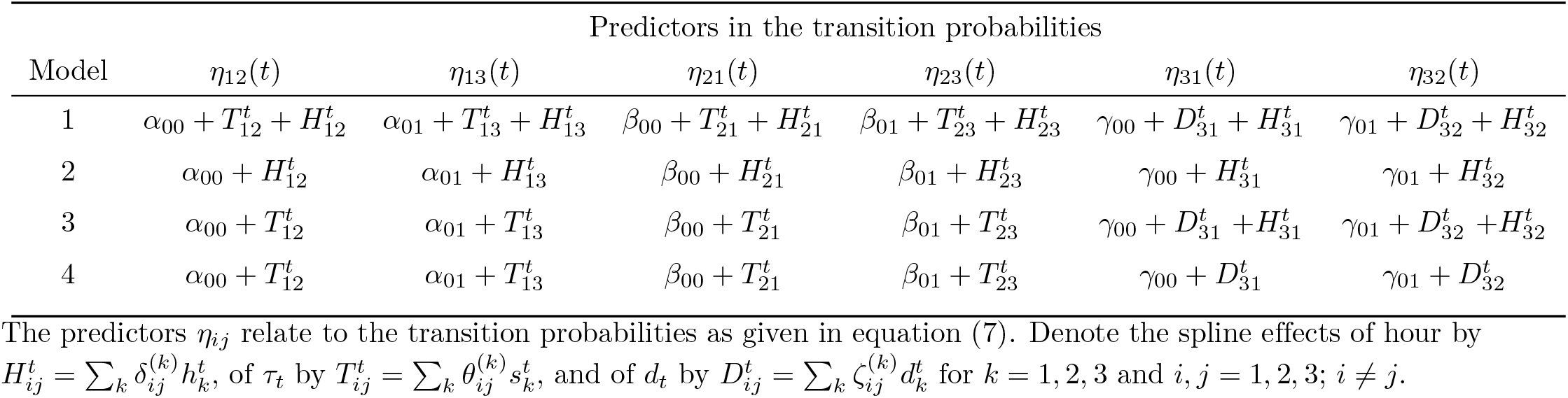
Different models for covariate effects on the transition probabilities between behavioural states.

### 2.4 The likelihood function and optimization

The likelihood *L_T_* of *x*_1_, *x*_2_,…, *x_T_*, where *x_t_* is the observation of ***X**_t_*, assumed to be generated by an *m*-state HMM, can in general be computed recursively in only *O*(*Tm*^2^) operations by the forward algorithm [22]. The likelihood is expressed as

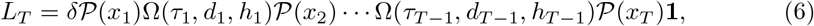

where 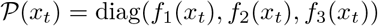 is a diagonal matrix with diagonal elements *f_i_*(*x_t_*) given in Eq. (4) when the gamma distribution is used, or Eq. (5) when the log-normal distribution is used, Ω is given by Eq. (1) and 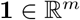 is a column vector of ones. The initial state distribution is denoted by *δ*, which is an *m*-dimensional row vector; *δ_i_* = *P*(*C*_1_ = *i*). For *δ*, we choose the uniform distribution, *δ_i_* = 1/*m*. Alternatively, it can be estimated, but there is no need for this extra computational effort, since our dataset is large and the influence of *δ* will be negligible. Furthermore, *δ* has no particular biological relevance.

The transition parameters in Eq. (1) are constrained to be between 0 and 1 with row sums equal to 1, and thus, even if there are 3^2^ = 9 entries, there are only 3 · 2 = 6 free parameters. To obtain an unconstrained optimization problem, we reparametrise to working parameters, as also done in [5,16,19], see also [22], by defining

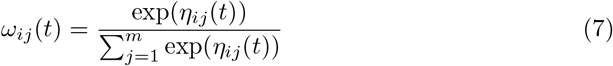

where *η_ij_* (*t*) is the predictor for dive *t* for 1 ≤ *i,j* ≤ 3, *i* ≠ *j*, and *η_ii_* = 0 for *i* = 1, 2, 3. This assures positive entries and that rows sum to 1.

We used the direct numerical Newton-Raphson algorithm `nlm` (`optim` in case `nlm` failed) in R [20] to estimate the parameters of the model by maximizing the log-likelihood, 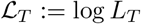, where *L_T_* is given in Eq. (6). The procedure `ns` from the package `splines` (version 3.5.0) was used to calculate the periodic splines. The procedure `pbs` from the package `pbs` (version 1.1) was used to calculate the periodic splines.

Using a combination of R and Rcpp [6] for calculating the log-likelihood function 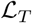 improved the runtime considerably. To mitigate the problem of local maxima, we ran the optimization algorithm thousands times with different starting values for the parameters of the state-dependent distributions, as well as for the parameters of the transition matrix. The final result was chosen as the one giving the maximum log-likelihood. For the best fits, we employed the jittering procedure used in [5], but it did not improve the log-likelihood value.

Once the optimal model was selected and parameters of the model were estimated, it was of interest to decode the most likely state sequence 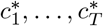. The Viterbi algorithm [7,22] is suitable to estimate the hidden states given the observed depths and durations:

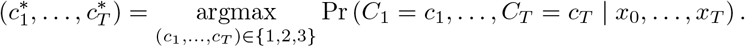

We evaluate by AIC which model provides the best fit, and study if final conclusions on diving behaviour differ if dependence is assumed.

## 3 Results

The estimation algorithm classifies each dive to one of the three hidden states. The classification depends on the model, but all models roughly group dives according to maximum depth. The algorithm allocates labels arbitrarily, so to compare across models we relabeled the states, such that State 1 represents the shortest and shallowest dives, which we define as near-surface travelling, State 2 represents medium long and deep dives, which we define as a feeding state, and State 3 represents the deepest and longest dives, which we define as an intense deep feeding state. Thus, one goal of comparing models is to access if conclusions on diving behaviour expressed through the decoded classes of the dives differ between models. If they all classify the same, it does not matter which model we use, maybe except for the estimation of covariate effects. If the classification differ from model to model, it is important to choose the statistically best model.

Table 2 lists the model selection results from the optimization. We use AIC to select the best model, which is highlighted in bold. The correlated log-normal model is clearly preferred above the independent models, with huge AIC differences, and the log-normal distribution is clearly preferred above the gamma distribution. Models with ΔAIC larger than 10 have essentially no support in the data compared to the best model [2]. Model 1 is the best among the tested models for all state distribution models, which balance accuracy and complexity of the model. It has diurnal effects on all transition probabilities, and nonlinear effects of *τ_t_* and *d_t_* on some of the transition probabilities. The marginal fit is illustrated in Fig 5 for correlated log-normal state distributions. The fits look convincing.

**Table 2.** Model selection results.

|  | Independent<br>Gamma<br>distribution |  | Independent<br>Log-normal<br>distribution |  | **Correlated**<br>**Log-normal**<br>**distribution** |  |
| --- | --- | --- | --- | --- | --- | --- |
| Model | $np$ | $\Delta AIC$ | $np$ | $\Delta AIC$ | $np$ | $\Delta AIC$ |
| **1** | 60 | 5050.51 | 60 | 2309.12 | **63** | **0** |
| 2 | 42 | 5386.94 | 42 | 2652.4 | 45 | 255.97 |
| 3 | 48 | 5096.47 | 48 | 2353.17 | 51 | 34.34 |
| 4 | 42 | 5194.44 | 42 | 2451.91 | 45 | 166.88 |

Differences in AIC values, $\Delta AIC$ , between the different models, where all AIC values were subtracted the AIC value of the model with the lowest AIC. For all the tested state distributions, covariate model 1 was preferred. The best fit is given by the minimum AIC, which occurs for the dependent log-normal state distribution, highlighted in bold. $np$ : number of parameters.

**Fig. 5.**
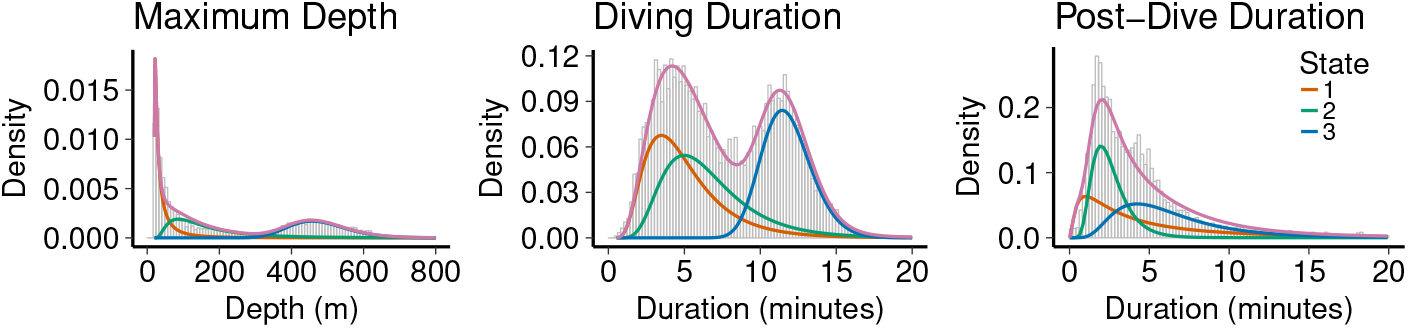
Model fit. Histograms of response variables Maximum Depth, Duration of Dives and Post-dive Duration. The fit of Model 1 with correlated log-normal distributions is indicated with a magenta curve, and the distribution of the fitted states are indicated with colours as given in the legend. State 1 corresponds to near-surface travelling, state 2 is feeding at medium depths, and state 3 is intense feeding at large depths.

A deep dive is defined to be a dive of at least 350 *m*, and therefore *d_t_* will typically be 0 once the narwhal is in state 1 or 2, and thus, we only expect *d_t_* to influence *η*_3*j*_. This is exactly what we see; models with no dependence of *d_t_* in transitions from states 1 and 2 are preferred over the other models (results not shown). Furthermore, we would expect that low or moderate values of *d_t_* would make the propensity to stay in state 3 large, and thus, decrease the probabilities for transiting to other states, whereas large values of *d_t_* are expected to increase the probabilities of changing, because the narwhal might need to rest. Thus, a non-linear relation might be adequate. This is explored in Models 1, 3 and 4, and effectively, the optimal model 1 does include a non-linear relationship of *d_t_*. Fig 6 illustrates the estimated covariate effects, and parameter estimates and confidence intervals can be found in Tables 4 and 5 in the Supplementary Material.

**Fig. 6.**
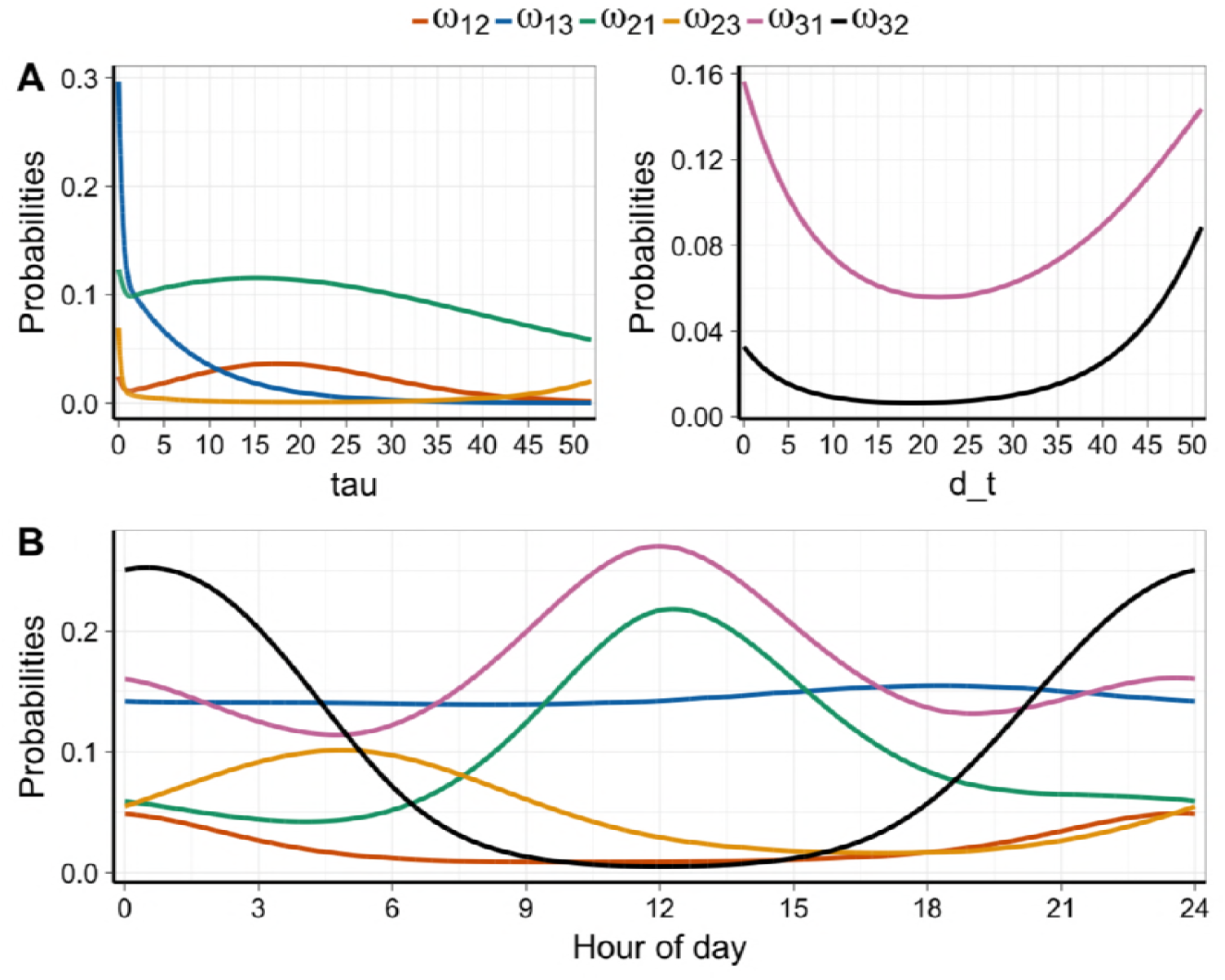
Covariate effects. A: Transition probabilities between behavioural states depending on covariates of correlated log-normal model 1, at approximately 12 pm. B: Transition probabilities depending on diurnal effects in model 1 with correlated log-normal state distributions, calculated for *τ_t_* = 0.58 and *d_t_* = 0 (the medians).

The covariate *τ_t_* indicates the time passed since last deep dive. We expect that *τ_t_* has impacts on states 1 and 2, but not on state 3 (which is the case for the selected model). In the left panel of Fig 6A the effect of *τ_t_* is illustrated. The transition probabilities do not seem to depend much on *τ_t_*, except for the probability of changing from surface travelling to deep dives. The probability is much higher for very small values of *τ_t_*, and decreasing fast towards 0 for larger values. This is not what was expected, but might reflect the following. When short time has passed since last deep dive, it is probably also short time since the whale was in state 3 of intense feeding. Thus, it reflects that the whale is still in an overall behavioral state of intense feeding, but just had a short break with surface travelling. This phenomenon can be seen in Fig 7 where the state decoding is shown for 12 representative hours. It is seen that after (at least) six dives in state 3, the whale changes to a few shallow dives for a short time, and then continues with another three dives in state 3. When a little longer time passes, the whale has effectively stopped its intense foraging, and the probability of a change to state 3 becomes smaller. Then, when long time has passed, we expect the transition probability to increase, which is not what is estimated. However, there are very few large observations of *τ_t_*: 75% of the values are belov 2.8 hours, and 90% are below 7.8 hours. Therefore effect estimates for large values are unreliable. The same is true for covariate *d_t_*: more than half are 0, 75% are 2 or smaller, and 90% are 8 or lower.

**Fig. 7.**
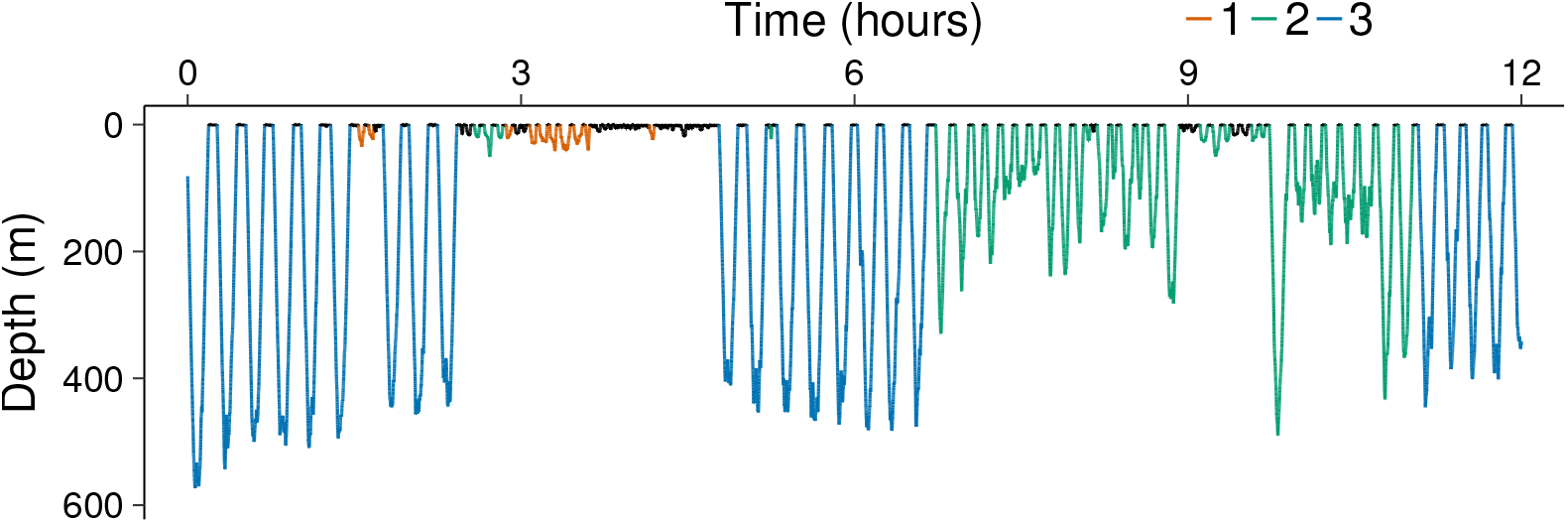
State decoding close-up. The estimated hidden state per dive for 12 hours of the data.

The effect of *d_t_* is illustrated in the right panel of Fig 6A. As expected, for values above 20 dives in a row, the probabilities to exit state 3 increase with increasing *d_t_*. The probability of changing to shallow waters (state 1) is much higher than the probability of changing to medium deep dives (state 2) after a period of intense feeding (state 3).

Fig 6B shows the diurnal effects on the transition probabilities. Changing from state 3 to 2 has highest probability around midnight, whereas changing from state 2 to 3 has highest probability around 6 pm. Changing to state 1 has highest probability around noon. The transition probabilities from state 1 do not depend much on diurnal effects.

Table 3 lists the estimated means and standard deviations of the state distributions for correlated log-normal and independent gamma distributions, respectively. Means and standard deviations of maximum depth and diving time are estimated larger for both shallow dives (state 1) and medium dives (state 2) with the correlated model compared to the independent model, whereas the two models estimate mean and variances approximately the same for deep dives (state 3). Thus, taking into account the dependence between the two state variables reveals more variable diving patterns (i.e., larger variance within states), unless the narwhal is doing intense feeding at deep dives, where the need for regular breathing do not allow the whale to digress.

**Table 3.** Summary measures of Model 1.

|  | State 1 | State 2 | State 3 |
| --- | --- | --- | --- |
| **Correlated Log-normal distribution** |  |  |  |
| $E_{MD}$ | 34.40 | 160.44 | 459.41 |
| $SD_{MD}$ | 79.66 | 143.22 | 83.14 |
| $E_{DT}$ | 5.08 | 6.63 | 11.78 |
| $SD_{DT}$ | 2.71 | 3.02 | 1.65 |
| $E_{PD}$ | 6.65 | 2.58 | 6.49 |
| $SD_{PD}$ | 10.61 | 1.16 | 3.73 |
| Corr | 0.56 | 0.81 | 0.46 |
| **Independent Log-normal distribution** |  |  |  |
| $E_{MD}$ | 25.95 | 135.55 | 458.00 |
| $SD_{MD}$ | 52.94 | 123.59 | 85.86 |
| $E_{DT}$ | 4.39 | 7.01 | 11.81 |
| $SD_{DT}$ | 2.16 | 2.20 | 1.67 |
| $E_{PD}$ | 6.85 | 2.65 | 6.64 |
| $SD_{PD}$ | 11.59 | 1.13 | 4.18 |
| **Independent Gamma distribution** |  |  |  |
| $E_{MD}$ | 20.46 | 113.71 | 455.46 |
| $SD_{MD}$ | 22.09 | 95.05 | 87.47 |
| $E_{DT}$ | 4.10 | 6.84 | 11.78 |
| $SD_{DT}$ | 1.75 | 2.21 | 1.70 |
| $E_{PD}$ | 8.24 | 2.56 | 7.32 |
| $SD_{PD}$ | 9.5 | 1.03 | 5.09 |

Means and standard deviations of Model 1 based on correlated Log-normal, independent Log-normal and independent Gamma distribution. MD: Maximum Depth; DT: Diving Time; PD: Post-Dive duration. E: mean; SD: standard deviation; Corr: Correlation.

Fig 8 shows the decoded hidden states for Model 1 with dependent log-normal state distribution. The correlated model estimates that the narwhal spends around 33.5% of its dives, corresponding to 34.4% of the time, in State 1 of near-surface travelling, which encompasses dives down to 793 *m* of durations up to 28 minutes. This is a large value for the surface state, but it is only the extreme tail of the distribution, and is represented by a single dive. It reflects that the log-normal distribution has heavier tails than the gamma distribution. Of the time spent in state 1, only 36.4% of the time is spent diving, the rest of the time the whale is in the surface. The narwhal spends around 32.5% of its dives, corresponding to 19.9% of the time, in medium depths of between 20 *m* and 836 *m* and durations between 0.7 and 22 minutes. Also here, a few deep dives are decoded as medium dives. Of the time spent in state 2, 69.9% of the time is spent diving, the rest of the time the whale is in the surface. Finally, 34.0% of dives, corresponding to 45.9% of the time, are spent during intense feeding at depths between 231.5 *m* and 910.5 *m* and durations between 7.2 and 19.5 minutes. Of the time spent in state 3, 62.8% of the time is spent diving, the rest of the time the whale is in the surface. Fig 7 illustrates a close-up of the decoding of dives for an example period of 12 hours. The correlated model thus decodes a few of the deep dives as pertaining to states 1 and 2, probably because of these dives taking longer time than the deep dives decoded as state 3.

**Fig. 8.**
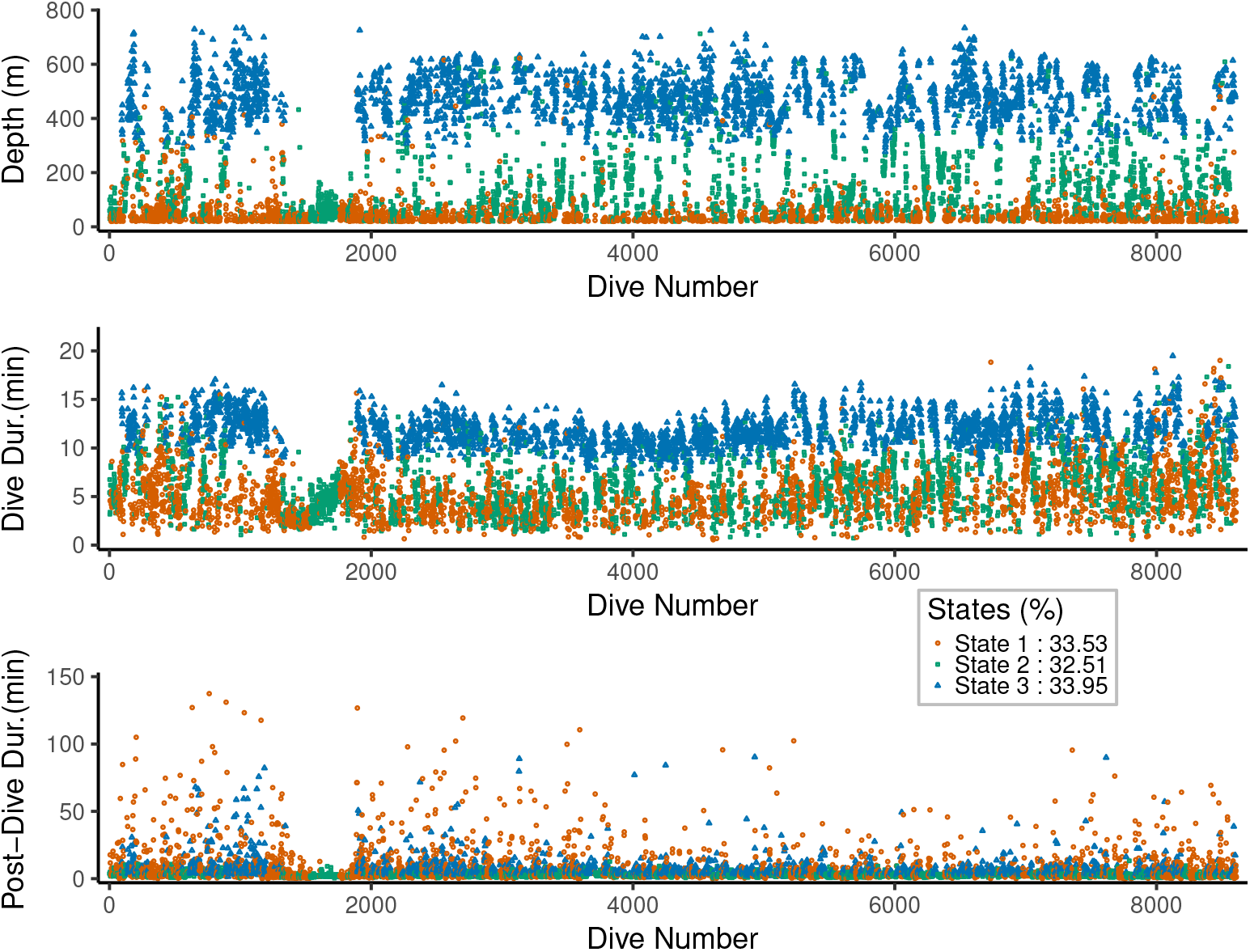
State decoding. The estimated hidden state per dive for each of the three observed variables under covariate model 1 and state distribution the correlated log-normal. The longest pause of no deep dives starts from the 1345th dive until the 1894th dive, and it lasts approximately 2 days and 17.5 hours.

Model 1 with an independent gamma state distribution estimates that the narwhal spends around 28.9% of its dives in State 1 of near-surface travelling, which encompasses dives up to 258 *m* of durations up to 14.8 minutes. It spends around 34.8% of its dives in medium depths of between 20 *m* and 584 *m* and durations between 1.5 and 28 minutes. Finally, 36.3% of dives are spent during intense feeding at depths between 172.5 *m* and 910.5 *m* and durations between 6.5 and 22 minutes. The independent model seems to be mostly guided by the maximum depth.

The decoding of behavioural states is thus different depending on whether correlation between observed variables are accounted for or not. The independent gamma model estimated that the narwhal spent 71.1% of its dives for feeding, whereas when correlation is taken into account, this estimate drops to 66.5%. These numbers are not too different, but to explore the differences further, we plot the distributions of the state variables and the covariates as a function of the decoding for the independent gamma and the correlated log-normal models in Fig 9. The orange density plots show the dives that are decoded the same in the two models, whereas the blue density plots show the dives that are differently decoded in the two models. In the correlated model, the distributions of dive characteristics in State 1 and 2 are broader, whereas they are more peaked in state 3, compared to the independent model.

**Fig. 9.**
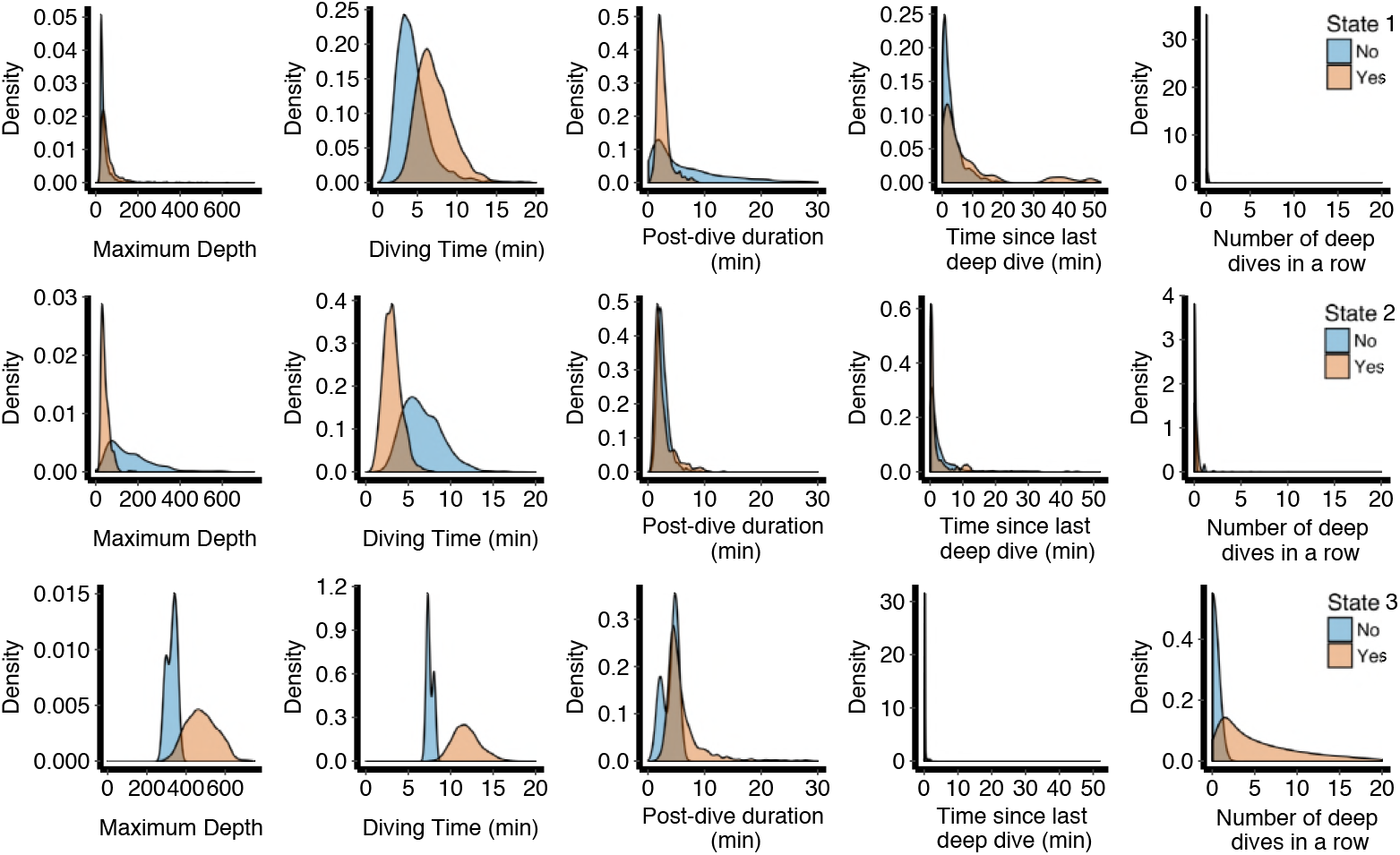
Distribution of hidden states. The empirical distributions of the response variables and the covariates stratified by decoded state in the correlated log-normal model 1. Within each panel, the orange density plots depict the distribution where the independent gamma and the correlated log-normal distributions agree on the decoded states, the blue density plots are those where the two distributions disagree.

To check the fit of the model more than what is given in Fig 5, we calculated the pseudo-residuals [22], qq-plots can be found in Fig 10. The fit is acceptable, maybe except for a too small lower tail for the Maximum Depth variable. This is probably due to the threshold of a depth of 20*m* in the definition of a dive.

**Fig. 10.**
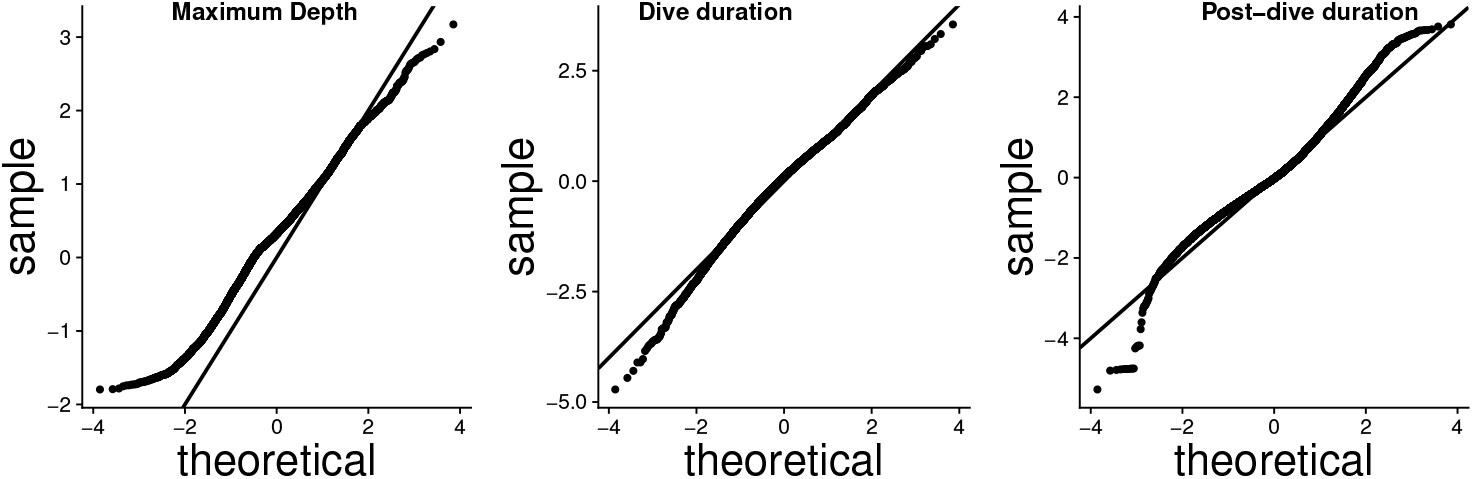
Quantile-quantile residual plots. QQ-plots of forecast pseudo-residuals from covariate model 1 with correlated log-normal state distribution.

## 4 Discussion

In this study, we investigate different multivariate HMMs with covariate effects for modelling the diving activity of a narwhal in the vertical dimension in the water column. The data set is extraordinarily long, which has made it possible to describe diurnal patterns in diving behaviour.

We extend the usual HMM models for diving behaviour of marine mammals to allow for dependence between state distributions, and show that the dependence has some impact on the conclusions drawn about the diving behaviour. We find that statistically the correlated model clearly outperforms the independent model, and more importantly, conclusions on the diving behaviour differ between the two models. The main differences are that the correlated model estimates more variable state distributions of maximum depth and dive duration, and that 66.5% of the dives are for feeding, compared to 71.1% in the independent model, under the assumption that states 2 and 3 in fact are representing feeding states in both models. Furthermore, the estimated means and variances of depth and duration of shallow and medium dives are estimated larger in the correlated model. Finally, ignoring the dependence between response variables usually leads to too narrow confidence intervals on parameter estimates.

Direct observations of feeding events were limited to the first week of the diving data but the depths where feeding events were detected served as a valid proxy for the depth threshold between behavioural state 2 and state 3. The observation that feeding events involve deep dives (≥ 350*m*) is also supported by studies of the buzzing activity during dives to different depths for narwhals travelling in the same area and time of the year as the whale included in this study [3].

Apparently the whale could stay in state 1 and 2 for long periods (> 24 hours) without transiting to state 3, and it even showed a pause of almost 3 days without deep dives. This indicates that feeding occur infrequently and that narwhals at least during summer and fall may have extended periods without feeding activity (see also [13]). However, the median of these pauses without state 3 dives was 44 minutes and the mean was 2 hours.

Transition from state 1 to presumed feeding activity is more likely to be to state 3 with deep dives, and rarely goes to state 2 from state 1. Diving activity in state 3 usually last for a series of dives (5-10) perhaps indicating that specific layers of prey is being detected and explored for a series of dives before the whale needs to spend an extended period at the surface. The post dive time is typically around 6.5 minutes after a state 3 dive, whereas it is typically only 2.6 minutes after a state 2 dive. The whale probably needs to spend more time at the surface to recover from nitrogen tissue tension following a longer breath-hold diving activity.

Even though detailed dive information supplemented by data on feeding events have been available for this analysis it may still not be adequate for describing the important drivers of diving behaviour. Both physiological constrains and reproductive state as well as environmental conditions may influence the diving activity to an extent that cannot be fully discerned in HMM analysis of dive series. For logistical reasons it is very difficult if not impossible to obtain information on all factors that affect the diving behaviour. However, the analysis of dive series provides a minimal insight into the integrated effect of the various factors driving the diving behaviour and the major advantage of the HMM analysis probably relies in the objective inter- and intra-specific comparison of diving activity.

## Acknowledgements

The study was supported by the Greenland Institute of Natural Resources; the Danish Cooperation for the Environment in the Arctic, the Carlsberg Foundation and University of Copenhagen Excellence Programme for Interdisciplinary Research.

## Supplementary Materials

**Table 4.** Estimates of the model parameters (1st part) in model 1 and their 95% confidence intervals for correlated Log-normal distribution.

| Correlated log-normal distribution |  |  |
| --- | --- | --- |
|  | Estimate | 95% CI |
| $\mu_1^{MD}$ | 2.61 | [2.56, 2.66] |
| $\mu_2^{MD}$ | 4.78 | [4.73, 4.84] |
| $\mu_3^{MD}$ | 6.11 | [6.11, 6.12] |
| $\sigma_1^{MD}$ | 1.36 | [1.33, 1.39] |
| $\sigma_2^{MD}$ | 0.77 | [0.72, 0.81] |
| $\sigma_3^{MD}$ | 0.18 | [0.17, 0.19] |
| $\mu_1^{DT}$ | 1.50 | [1.48, 1.52] |
| $\mu_2^{DT}$ | 1.80 | [1.77, 1.82] |
| $\mu_3^{DT}$ | 2.46 | [2.45, 2.46] |
| $\sigma_1^{DT}$ | 0.50 | [0.49, 0.51] |
| $\sigma_2^{DT}$ | 0.43 | [0.41, 0.46] |
| $\sigma_3^{DT}$ | 0.14 | [0.14, 0.14] |
| $\mu_1^{PD}$ | 1.26 | [1.22, 1.3] |
| $\mu_2^{PD}$ | 0.86 | [0.83, 0.88] |
| $\mu_3^{PD}$ | 1.73 | [1.71, 1.75] |
| $\sigma_1^{PD}$ | 1.13 | [1.10, 1.15] |
| $\sigma_2^{PD}$ | 0.43 | [0.41, 0.45] |
| $\sigma_3^{PD}$ | 0.53 | [0.52, 0.55] |
| $\rho_1$ | 0.56 | [0.53, 0.58] |
| $\rho_2$ | 0.81 | [0.78, 0.83] |
| $\rho_3$ | 0.46 | [0.43, 0.50] |

In state $i$ , $\mu_i$ and $\sigma_i$ are the log-mean and log-standard deviation of the correlated log-normal distribution. Index MD stands for Maximum Depth, DT stands for Dive Duration and PD stands for Post-Dive time. The depth is measured in meters, and time in seconds. The confidence intervals were computed from the Hessian of the negative log-likelihood function, i.e., based on the inverse of the observed Fisher information (to be continued on next page).

**Table 5.**
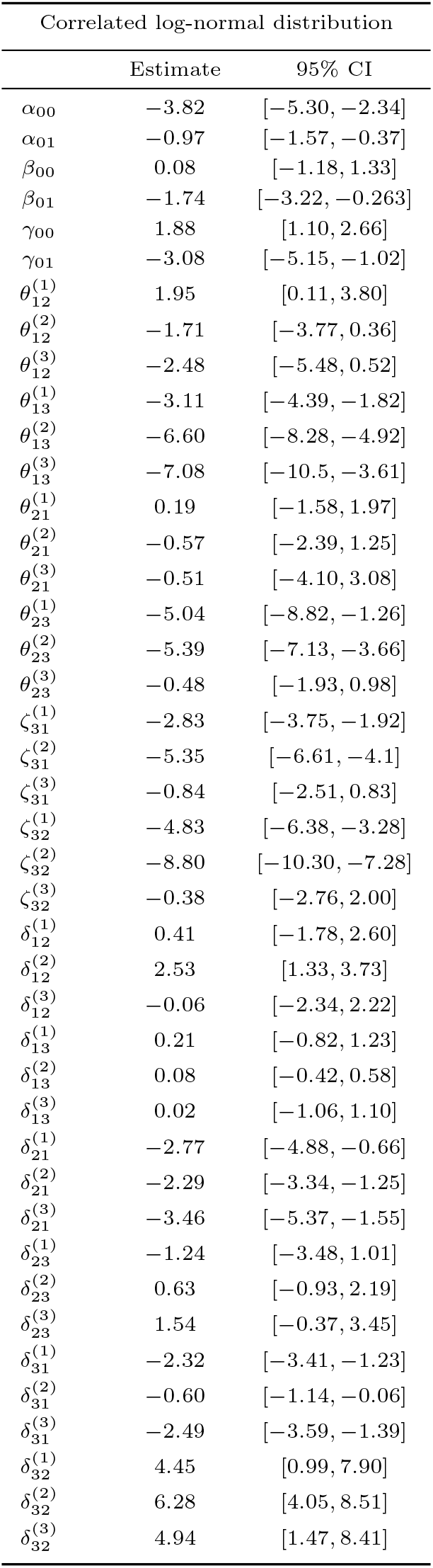
Estimates of the model parameters (2nd part).

